# Semaglutide promotes intramuscular fat formation after injury

**DOI:** 10.64898/2026.06.16.732451

**Authors:** Christian Noble, Dawson Geller, Nikhil Urs, Daniel Kopinke

## Abstract

Glucagon-like peptide 1 receptor agonists (GLP-1RAs) have become defining therapies in the management of type 2 diabetes and obesity. Despite recent interest in the effects of GLP-1RA therapy on skeletal muscle, their influence on muscle repair after injury remains largely untested. Because GLP-1RA use is common in populations at heightened risk for diminished regenerative capacity, a critical unanswered question is whether GLP-1R agonism supports muscle regeneration or alters the normal course of recovery after injury. Using intramuscular glycerol injection as an adipogenic injury model, we assessed whether semaglutide, a widely prescribed GLP-1RA, alters the balance between myogenesis and adipogenesis during regeneration. Surprisingly, semaglutide treatment markedly increased the formation of intramuscular adipose tissue (IMAT) and inhibited the growth of regenerated fibers. These effects were injury-dependent, as uninjured muscle showed no detectable differences in IMAT or myofiber size. Together, these findings identify a previously underappreciated context in which GLP-1RA therapy may adversely affect muscle quality.

## Main text

Glucagon-like peptide 1 receptor agonists (GLP-1RAs) have become defining therapies in the management of type 2 diabetes and obesity. With their widespread adoption, concerns regarding the substantial lean mass loss reported by clinical trials have inspired recent investigation into the effects of GLP-1RA therapy on skeletal muscle size and strength^1,2^.However, their influence on muscle repair after injury remains largely untested. Because GLP-1RA use is common in populations at heightened risk for diminished regenerative capacity, such as patients with diabetes and older adults^3,4^, a critical unanswered question is whether GLP-1R agonism supports muscle regeneration or alters the normal course of recovery after injury.

Effective muscle regeneration requires precise coordination of muscle stem cells to repair damaged myofibers. This process relies on pro-regenerative signaling from fibro-adipogenic progenitors (FAPs), a muscle-resident mesenchymal stromal population activated by injury. When regeneration is disrupted, contractile tissue is replaced by fibrosis and intramuscular adipose tissue (IMAT), the pathological accumulation of adipocytes between myofibers driven by aberrant FAP differentiation. IMAT is strongly associated with poor muscle quality and insulin resistance in diabetes, obesity, and aging and is increasingly recognized as a contributor, not merely a correlate, of impaired muscle function^5–7^.

Here, we asked a straightforward question: does semaglutide, a widely prescribed GLP-1RA, alter the balance between myogenesis and adipogenesis during regeneration? To answer this, we injured the tibialis anterior (TA) muscles of adult C57BL/6J mice (n = 40 total; ∼equal females and males) by intramuscular glycerol injection, a commonly used injury model that reliably induces adipogenic remodeling. Beginning immediately after injury, mice received daily intraperitoneal injections of semaglutide (50 µg/kg) or vehicle for two weeks (Fig 1A). EchoMRI body composition measurements collected before and after treatment confirmed expected systemic effects consistent with known GLP-1RA-driven weight reduction. Semaglutide reduced total body mass (∼6% in females and 12% in males) with decreases in both fat mass (∼27% and ∼50%) and lean mass (∼3% and 7%) (Fig 1B-1D).

**Figure 1.**
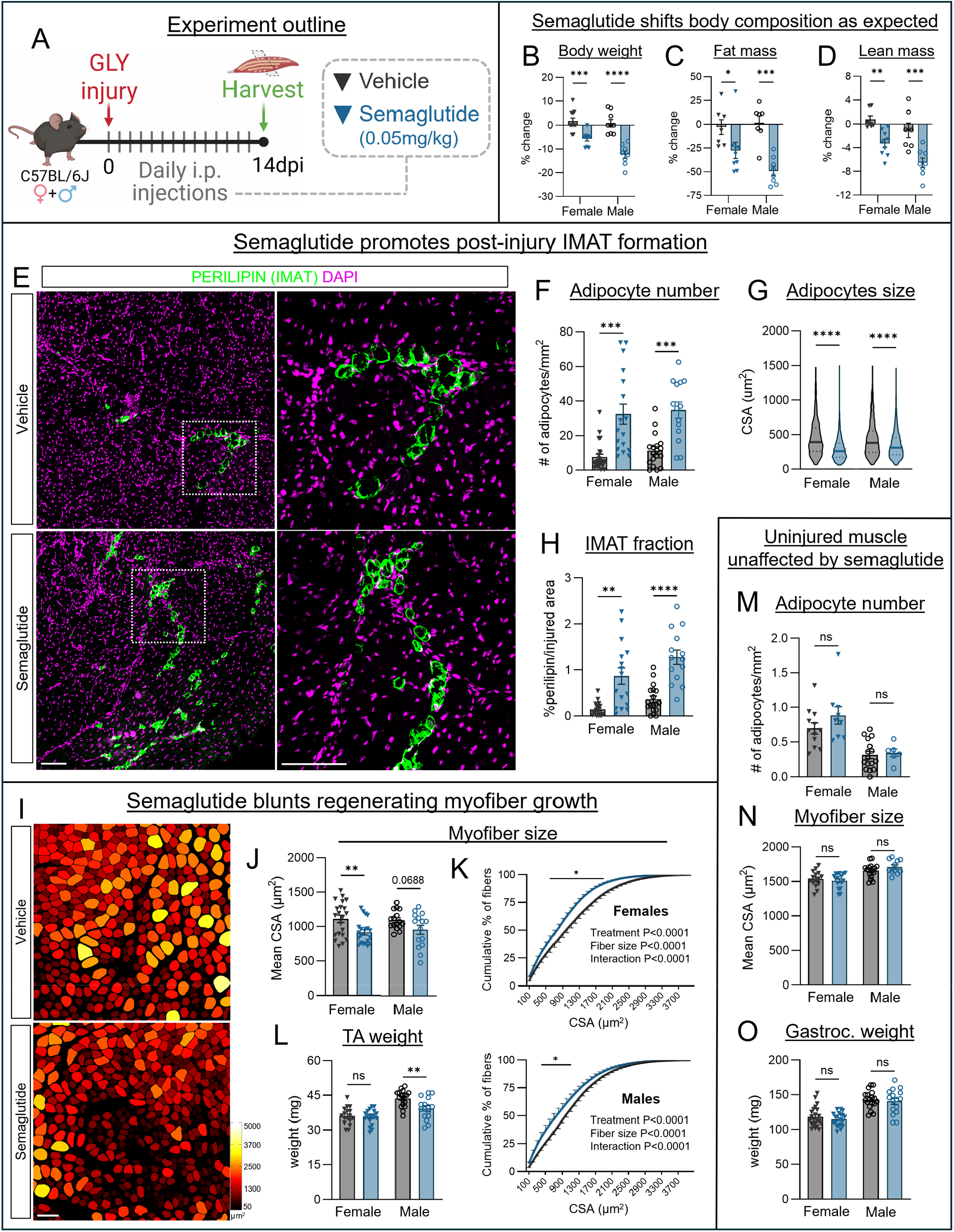
(A) Experiment timeline. Following 14 days of semaglutide or vehicle injections: (B) Percent change in body weight from baseline. Percent change in fat mass (C) and lean mass (D) from baseline, measured using EchoMRI. (E) Immunofluorescence images of transverse sections of tibialis anterior muscles from vehicle- and semaglutide-treated mice depicting PERILIPIN^+^ adipocytes (green) and DAPI (magenta). (F) Number of adipocytes per mm^2^ of injured muscle. (G) Adipocyte cross-sectional area (CSA). (H) Percentage of perilipin-positive area relative to total injured muscle area. (I) Transverse sections of tibialis anterior muscles depicting regenerated myofibers pseudo-colored by CSA. (J) CSA of regenerated myofibers. (K) Cumulative distribution of myofiber CSA in females (top) and males (bottom). (L) Tibialis anterior weight at harvest. Uninjured gastrocnemius muscles: (M) number of adipocytes per mm^2^, (N) myofiber CSA, (O) muscle weight at harvest. Representative images are from female mice. All scale bars: 100 μm.

Surprisingly, histological analysis of injured TA muscles revealed that semaglutide shifted regenerative outcomes toward pathological adipogenic remodeling. Semaglutide-treated mice exhibited a robust increase in PERILIPIN^+^ adipocytes interspersed among regenerating myofibers, with ∼5-fold greater adipocyte number relative to vehicle controls (Fig 1F). Consistent with the clinical fat-shrinking effect of GLP-1R therapy, individual IMAT adipocytes were smaller on average in semaglutide-treated muscles (Fig 1G). Despite this apparent “anti-hypertrophic” effect on adipocyte size, the net consequence was a markedly larger IMAT burden, as the fraction of injured muscle area occupied by IMAT increased ∼4-fold compared with controls (Fig 1H).

In parallel, semaglutide appeared to delay or inhibit myofiber regeneration. Quantification of nascent, centrally-nucleated myofibers demonstrated reduced cross-sectional area (CSA) in semaglutide-treated mice. On average, regenerated fibers were smaller in females, with males approaching significance (Fig 1J). Cumulative frequency distributions of nascent myofiber CSA revealed a leftward shift toward smaller fibers in both sexes (Fig 1K). These findings show that semaglutide impaired the normal restoration of myofiber size during regeneration, which is indicative of a detrimental effect on muscle recovery and is consistent with the recent finding of another group^8^. In males, this occurred alongside a reduction in TA weight (∼10%) (Fig 1L).

Analyses of uninjured gastrocnemius muscles support the interpretation that semaglutide’s observed effects on skeletal muscle are context-dependent and occur specifically within injured, regenerating muscle. Gastrocnemius muscles displayed little to no IMAT (Fig 1M), as expected for uninjured murine muscle^6^, and there were no detectable differences in weight (Fig 1O) or myofiber CSA (Fig 1N) between treatment groups in either sex. This aligns with a prior study suggesting that the magnitude of semaglutide-associated lean mass loss is not reflected in loss of hindlimb muscle weight or myofiber size^1^.

Taken together, greater IMAT deposition and impaired regenerative hypertrophy of myofibers represent a clear sign of compromised regeneration and diminished muscle quality. This observation is clinically provocative because it runs counter to the intuitive expectation that an anti-obesity therapy would lessen fat accumulation in peripheral tissues. Instead, we found that semaglutide promoted intramuscular adipocyte formation after injury even as whole-body adiposity declined, suggesting that local adipogenic remodeling in injured muscle can be enhanced during GLP-1RA therapy. Importantly, MRI studies of muscle composition have largely focused on uninjured muscle conditions, where incretin-based therapy has been reported to modestly reduce thigh muscle fat infiltration in people with type 2 diabetes^9,10^. However, MRI-derived fat infiltration typically reflects total lipid content and is unable to distinguish IMAT in particular. These studies are often cited as evidence of favorable effects on muscle quality, but our findings bring to light the unconsidered yet critical context of acute muscle damage and repair, highlighting the need to evaluate GLP-1R agonism during periods when muscle is actively remodeling. Notably, because commonly used mouse strains, particularly C57BL/6J, are resistant to IMAT formation^6^, the robust increase in intramuscular adipocyte number we observe may underestimate the potential magnitude of this effect in humans, who are generally more prone to form IMAT^6^ . This may be especially relevant in clinical settings where muscle is critically compromised and susceptible to fatty degeneration such as in chronic limb-threatening ischemia (CLTI), the most severe form of peripheral artery disease (PAD). The STRIDE trial recently showed that semaglutide improved walking capacity in patients with mild PAD and type 2 diabetes^11^. However, it excluded patients with more advanced stages of the disease such as CLTI, which is distinguished by extensive fibrofatty degeneration of skeletal muscle with IMAT content being a salient predictor of impaired limb function^12^. Thus, it will be important to determine whether GLP-1R agonism remains beneficial when muscle is under ischemic and adipogenic stress.

This short study was not designed to define mechanism, but the results support several testable explanations. First, since semaglutide reduces energy intake and body mass, caloric restriction during regeneration could blunt anabolic signaling required for myofiber growth. Second, the concomitant increase in IMAT could impede regeneration via physical constraints on nascent myofibers. Third, and most intriguingly, the IMAT phenotype itself being driven by enhanced adipogenesis, rather than adipocyte hypertrophy, points to FAP fate regulation as a likely site of action. To our knowledge, direct effects of GLP-1RA therapy on FAP proliferation, survival, or adipogenic differentiation have not been characterized. Whether semaglutide acts cell-autonomously (via GLP-1R-dependent signaling in FAPs) or indirectly by altering the post-injury environment (e.g. immune response and nutrient availability) remains an important open question. Moreover, we intentionally used metabolically healthy mice to isolate semaglutide’s influence on muscle regeneration from the confounding effects of pre-existing obesity (driven genetically or by high-fat diet), which independently impair regeneration and promote adipogenic remodeling^3^. Dedicated studies in models of obesity or diabetes will be important to determine whether these effects are amplified or altered when the regenerative niche is already biased toward adipogenesis.

In summary, semaglutide treatment during the acute regenerative window unexpectedly increased IMAT formation and reduced regenerating myofiber size after glycerol injury. The absence of detectable differences in uninjured muscle over the same period support that the effects of GLP-1R agonism are injury-dependent. These results identify a previously unrecognized potential risk of GLP-1RA therapy for the muscle health of patients. We hope this report stimulates mechanistic studies to define how GLP-1RA treatment influences the recovery trajectory of injured muscle and encourages greater incorporation of muscle quality and regeneration-focused outcomes into future clinical investigations.

## Methods

### Muscle injury and treatment

All injuries were performed on adult (11–14-week-old) C57BL/6J mice. Mice were put under anesthesia with isoflurane, and the tibialis anterior (TA) muscles were injected with 30–50 μL of 50% glycerol (GLY; Acros Organics, 56-81-5) diluted in sterile saline. Injuries were performed bilaterally. Beginning at the time of injury, experimental mice received daily intraperitoneal injections of semaglutide (AdipoGen Life Sciences; Cat# AG-CP3-0040-M025) at a dose of 50μg/kg, while control mice received vehicle at equal volume. Mice were housed under a 12-hour light/dark cycle with *ad libitum* access to food and water. At 14 days post-injury, mice were euthanized by isoflurane inhalation, and tissues were weighed immediately at the time of harvest. Body composition was assessed using nuclear magnetic resonance (EchoMRI™-700 Analyzer) before the treatment period and at 11 days post-injury. All animal work was approved by the Institutional Animal Care and Use Committee (IACUC) of the University of Florida.

### Histology and immunofluorescence

Tibialis anterior (TA) and gastrocnemius muscles were fixed in 4% paraformaldehyde (PFA) for 2.5 h at 4°C, washed, and cryoprotected overnight in 30% sucrose at 4°C. Muscles were then embedded in OCT-filled cryomolds (Sakura Finetek; Cat# 4566) and snap-frozen in liquid nitrogen-cooled isopentane. Cryosections (12 μm) were collected every 250-350 µm using a Leica cryostat. For immunofluorescence, sections were incubated in blocking solution (5% donkey solution in PBS containing 0.3% Triton X-100) at 4°C for 45 min at room temperature. Primary antibody anti-PERILIPIN (Cell Signaling Technology; Cat# 9349; 1:1000) in blocking solution was applied overnight at 4°C. Species-appropriate Alexa Fluor secondary antibodies (Thermo Fisher Scientific; 1:1000), DAPI (1 µg/mL), and Phalloidin-Alexa Fluor 647 (Thermo Fisher Scientific; Cat# A22287; 1:500) in blocking solution were applied for 1 hour at room temperature. Slides were mounted with Fluoromount-G (SouthernBiotech; Cat# 0100-01). Images were acquired on a Leica DMi8 using a 20x objective.

### Image analysis

To quantify myofiber cross-sectional area, images of sections stained against PHALLOIDIN were segmented through Cellpose and then processed through the ImageJ plug-in LabelsToRois. To assess regeneration in TA muscles, uninjured regions were excluded prior to analysis and were identified as areas containing myofibers without centrally located nuclei. PERILIPIN+ adipocytes were manually counted from tile-scan images of full muscle cross sections and normalized to injured area in TAs and total muscle area in gastrocnemius. The cross-sectional areas of intramuscular adipocytes were manually measured. IMAT fraction was calculated as PERILIPIN+ area divided by total injured muscle area. All images were processed and quantified through ImageJ Software (v1.552p).

### Statistical analysis

Data were graphed using GraphPad Prism (version 10) and are presented as mean ± SEM. For myofiber size distributions, two-way analysis of variance (ANOVA) was performed followed by Šidák’s multiple comparisons for pairwise comparisons. All other comparisons were performed using an unpaired two-tailed Student’s t test. Females and males were analyzed separately. A p-value of less than 0.05 was considered statistically significant, denoted as *p < 0.05, **p < 0.01, ***p < 0.001, and ****p < 0.0001.

